# Single cell RNA sequencing of 13 human tissues identify cell types and receptors of human coronaviruses

**DOI:** 10.1101/2020.02.16.951913

**Authors:** Furong Qi, Shen Qian, Shuye Zhang, Zheng Zhang

## Abstract

The new coronavirus (2019-nCoV) outbreak from December 2019 in Wuhan, Hubei, China, has been declared a global public health emergency. Angiotensin I converting enzyme 2 (ACE2), is the host receptor by 2019-nCov to infect human cells. Although ACE2 is reported to be expressed in lung, liver, stomach, ileum, kidney and colon, its expressing levels are rather low, especially in the lung. 2019-nCoV may use co-receptors/auxiliary proteins as ACE2 partner to facilitate the virus entry. To identify the potential candidates, we explored the single cell gene expression atlas including 119 cell types of 13 human tissues and analyzed the single cell co-expression spectrum of 51 reported RNA virus receptors and 400 other membrane proteins. Consistent with other recent reports, we confirmed that ACE2 was mainly expressed in lung AT2, liver cholangiocyte, colon colonocytes, esophagus keratinocytes, ileum ECs, rectum ECs, stomach epithelial cells, and kidney proximal tubules. Intriguingly, we found that the candidate co-receptors, manifesting the most similar expression patterns with ACE2 across 13 human tissues, are all peptidases, including ANPEP, DPP4 and ENPEP. Among them, ANPEP and DPP4 are the known receptors for human CoVs, suggesting ENPEP as another potential receptor for human CoVs. We also conducted “CellPhoneDB” analysis to understand the cell crosstalk between CoV-targets and their surrounding cells across different tissues. We found that macrophages frequently communicate with the CoVs targets through chemokine and phagocytosis signaling, highlighting the importance of tissue macrophages in immune defense and immune pathogenesis.

## Introduction

In December, 2019, a novel coronavirus (2019-nCoV) infection emerged in Wuhan. Over 30 thousand of people are infected with 2019-nCoV until February 8, showing that 2019-nCoV is highly contagious. Coronavirus is a type of single-stranded RNA (ssRNA) virus [1] Before the emergence of 2019-nCoV, there are 6 known human coronaviruses, including the Middle East respiratory syndrom coronavirus (MERS-CoV) and severe acute respiratory syndrome coronavirus (SARS-CoV). The symptoms caused by 2019-nCoV infection include acute respiratory distress syndrome (~29%), acute cardiac injury (~12%) or acute kidney injury (~7%)[2], implying that 2019-nCoV may infect various human tissues.

Viruses bind to host receptors on target cell surface to establish infection. Membrane proteins mediated membrane fusion allowed the entry of enveloped viruses[3]. As recently reported, both nCoV and SARS-CoV could use ACE2 protein to gain entry into the cells[4]. Since the outbreak, many data analysis have showed a wide distribution of ACE2 across human tissues, including lung[5], liver[6], stomach[7], ileum[7], colon[7] and kidney[8], indicating that 2019-nCoV may infect multiple organs. However, these data showed that AT2 cell (the main target cell of 2019-nCoV) in the lung actually expressed rather low levels of ACE2[8]. Hence, the nCoVs may depends on co-receptor or other auxiliary membrane proteins to facilitate its infection. It is reported that viruses tend to hijack co-expressed proteins as their host factors[9]. For example, Hoffmann et al. recently showed that 2019-nCoV-S uses ACE2 for entry and depends on the cellular protease TMPRSS2 for priming[10], showing that 2019-nCoV infections also require multiple factors. Understanding the receptors usage by the viruses could facilitate the development of intervention strategies. Therefore, identifying the potential co-receptors or auxiliary membrane proteins for 2019-nCoV is of great significance.

For this purpose, we collected single cell gene expression matrices from 13 relatively normal human tissues, consisting of lung[11], liver[12], ileum[13], rectum[13], blood[14], bone marrow[15], skin[16], spleen[17], esophagus[17], colon[18], eye[19], stomach[20] and kidney[21] from published literatures. We analyzed the single cell co-expression profiles of 51 known ssRNA viral receptors and 400 membrane proteins, including ACE2, in the identified 119 cell types across the 13 human tissues. Consistent with other recent studies, ACE2 was found to be present in lung, liver, stomach, ileum, kidney and colon. Using hierarchy clustering and correlation calculation, we found that ANPEP, DPP4 and ENPEP showed co-expression pattern with ACE2 across the 13 human tissues, suggesting that they may act as potential coreceptors for 2019-nCoV, and all human CoVs should have similar targets across different human tissues..After conducting “CellPhoneDB”, we further reveal that macrophages frequently crosstalks with CoVs-target cells, in multiple tissues.

## Materials and Methods

### Data collection

The gene raw counts or normalized gene expression matrix for each single cell were downloaded from GEO (https://www.ncbi.nlm.nih.gov/geo/) or Human Cell Atlas (https://www.humancellatlas.org) database (**Table S1**). In total, we collected single cell gene expression data of 13 tissues, including liver, lung, colon, ileum, rectum, blood, spleen, bone marrow, eye, skin, stomach, oesophagus and kidney. The data source and the sample information are listed as follows. Liver, GEO Accession No. GSE115469, 5 normal human donors; Lung, GEO Accession No. GSE130148, 4 human donors died from hypoxic brain damage; Colon, GEO Accession No. GSE116222, 3 healthy volunteers; Ileum and rectum, GEO Accession No. GSE125970, totally 4 intestine mucosae sampled at least 10 cm away from the tumor border; Skin, GEO Accession No. GSE132802, 4 healthy volunteers; PBMC, GEO Accession No. GSE136103, 4 samples from cirrhotic patients; Spleen and oesophagus, available from Human Cell Atlas, totally 11 cardiac death donors; Bone marrow, GEO Accession No. GSE120221, 5 healthy donors (A, E, J, R, U); Eye, GEO Accession No. GSE135922, 3 Macula and 3 periphery of human donor eyes; Stomach, GEO Accession No. GSE134520, 3 Non-atrophic gastritis patients; Kidney, GEO Accession No. GSE131685, 3 normal kidney tissues obtained at least 2 cm away from tumor tissue.

The high-quality virus-host receptor interactions were downloaded from Viral Receptor database (http://www.computationalbiology.cn:5000/viralReceptor), which curated 152 pairs of mammalian virus-host receptor interactions and 51 virus receptors from 9 mammal species. They curated five types of virus, consisting of single-stranded DNA (ssDNA) virus, doublestranded (dsDNA) virus, single-stranded (ssRNA) virus, double-stranded (dsRNA) virus and retro-transcribing virus. We only focus on ssRNA virus in this study. The membrane proteins were extracted from Membranome database (https://membranome.org).

### Data processing, quality control and normalization

The raw count matrix (UMI counts per gene per cell) was processed by Seurat[22]. Cells with less than 100 expressed genes (UMI count > 0) and higher than 25% mitochondrial genome transcript were removed. Genes expressed in less than three cells were removed. Then, we normalized the gene expression data using “NormalizeData” function with default settings. The sources of cell-cell variation driven by batch were regressed out using the number of detected UMI and mitochondrial gene expression, which was implemented by ‘‘ScaleData’’ function. The corrected expression matrix was used for cell clustering and dimensional reduction.

### Cell clustering, dimensional reduction and visualization

The cell clustering and dimensional reduction were performed by Seurat package. Before that, we choose 2000 highly variable genes (HVGs) from the corrected expression matrix and then centered and scaled them. It was implemented by ‘‘FindVariableGenes’’ function in the Seurat package. We then performed principle component analysis (PCA) on the HVGs using ‘‘RunPCA’’ function. To remove the signal-to-noise ratio, we select a number of significant principal components by implementing “JackStraw” function, which was implemented by permutation test. Specifically, we firstly identified 50 principal components as a result and then selected the significant components according to the p-values produced by “ScoreJackStraw” function for further analysis. The batch effects were removed by harmony package[23].

Cells were then clustered utilizing the ‘‘FindClusters’’ function through embedding cells into a graph structure in PCA space. We set the parameter resolution as 0.8 to identify only major cell types, *e.g*. T cells, B cells or macrophages. The clustered cells were then projected onto a two-dimensional space using “RunUMAP” function. The clustering results were visualized by “DimPlot” function.

### Cell type identification

To annotate cell clusters, we firstly identified the differentially expressed genes on each cluster by performing “FindMarkers” function. The cell clusters were then annotated according to curated known cell markers (**Figure S1**). The cell clusters consistently expressed the same cell marker were merged.

### Cell-cell interaction analysis

We conducted cell-cell interaction analysis utilizing cellphonedb function curated by CellPhoneDB database[24]. The significant cell-cell interactions were selected with p-value < 0.01.

## Results

### Cell type identification in 13 human tissues

We collected the single cell RNA sequencing data (raw count gene expression matrix or normalized gene expression matrix) from published literatures, which have been deposited in public database, *e.g*. GEO (https://www.ncbi.nlm.nih.gov/geo/) or Human Cell Atlas (https://www.humancellatlas.org). Totally, we curated single cell gene expression matrices of 13 human tissues, including lung[11], liver[12], ileum[13], rectum[13], blood[14], bone marrow[15], skin[16], spleen[17], esophagus[17], colon[18], eye[19], stomach[20] and kidney[21] (**Table S1**). The raw count matrix was normalized and scaled by Seurat package[22] (**Materials and Methods**). For each tissue, we performed cell clustering and dimension reduction on gene expression matrix using Seurat package. After filtering out low quality cells, we obtained 8443, 43474, 4248, 5282, 3279, 30693, 97695, 17131,4335, 11552, 4871, 8880, 20197 cells from liver, lung, colon, ileum, rectum, blood, spleen, bone marrow, eye, skin, stomach, esophagus and kidney, respectively (**Table S1**). The cell clusters were then annotated using canonical markers searched from the published articles (**Figure S1**). We finally annotated 119 cell types from 13 human tissues.

Lung belongs to respiratory system, in which 13 cell types were identified (**Figure S2**). These cell types consist of macrophages, Alveolar Type 2 cells (AT2), monocytes, NK&T cells, ciliated cells, basal cells, mast cells, neutrophils, Alveolar Type 1 cells (AT1), fibroblasts, endothelial cells, lymphatic cells and B cells. Among them, ~31% cells are alveolar cells (AT2 and AT1) and ~54% cells are immune cells (B cells, T cells and Myeloid cells).

Ileum, rectum, esophagus, colon, and stomach are part of digestive system. In esophagus, 7 cell types were identified (**Figure S2**). Keratinocytes show the highest percentage (~66%) of total cells. The remaining cells are B cells, epithelial basal cells, glands cells, stroma cells, T cells and vessel cells. We detected 11 cell types in stomach (**Figure S2**), in which epithelial cells (~29%) and pit mucous cell (PMCs) (~23%) constitute the largest group. Besides, B cells, endothelial cells (ECs), enteroendocrine cells, fibroblasts, antral basal gland mucous cell (GMCs), macrophages, neck-like cells, proliferative cell (PCs) and T cells were identified in stomach. In ileum (**Figure S2**), 7 cell types, including enterocytes (ECs), enteroendocrine cells (EECs), goblet cells (Gs), Paneth cells (PCs), progenitor cells (PROs), stem cells (SCs), transient amplifying (TAs), were annotated. Enterocytes are the largest cell population (~64%) in ileum. Rectum share cell types with ileum. Whereas, the largest cell population in rectum is progenitor cells (~37%). A total of 9 cell types was detected in colon (**Figure S2**). The percentage of colonocytes (colonocytes and crypt top colonocytes) is 53%, which constitute the largest cell population in colon. BEST4+ cells, enteroendocrine cells (EECs), goblet cells, innate lymphoid cells, mast cells, T cells and undifferentiated cells were also identified.

Liver, spleen and skin play vital roles in immune systems. Immune cells (~45%) and hepatocyte (~42%) account for the vast majority of cells in liver (**Figure S2**). Cholangiocytes, endothelial cells and erthyroid cells were also detected. Spleen is the immune organs in human body, in which all the cells are immune cells (**Figure S2**). Spleen composed of a large proportion of T cells (~33%) and B cells (~43%). Other immune cells, including CD34 progenitor cells, cDCs, dividing cells, innate lymphoid cells, macrophages, monocytes, neutrophils, NK cells and pDCs, also make up the spleen cell populations. Skin is a physical barrier against the external environment. We identified a total of 7 cell populations in skin (**Figure S2**), in which pericytes (~31%) and fibroblasts (~21%) are the most enriched populations. The immune cells, comprising T cells and myeloid cells were also identified. In addition, basal cells, endothelium cells, and suprabasal keratinocyte constitute the skin cell populations.

The kidneys are the part of urinary system that makes urine. Most of the cells in kidney are proximal tubule cells (Proximal Ts) (~82%) (**Figure S2**). Besides, we also annotated immune cells (~8%), collecting duct intercalated cells (Collecting DIs), collecting duct principal cells (Collecting DPs), distal tubule cells (Distal Ts) and glomerular parietal epithelial cells (Glomerular PEs) in kidney.

The eyes are sensory organs in nervous system. Ten cell types were identified in eyes (**Figure S2**). Fibroblasts and immune cells composed of ~31% and ~25% of total cells, respectively. Eyes also contain endothelial cells, melanocytes, pericytes, retinal pigment epithelium (RPEs) and Schwann cells.

Bone marrow is the primary site of hematopoiesis. We identified large number of NK/NKT cells (~44%) and erythrocytes (~28%) cells in bone marrow (**Figure S2**). B cells, hematopoietic stem cells, MK progenitors, monocytes, neutrophils and DCs were also detected in bone marrow.

Blood is circulated around various tissues. Monocytes (~32%) and T cells (~55%) make up the largest proportion of blood cells (**Figure S2**). In addition, we also identified B cells, cDCs, macrophages, NK cells, pDCs and platelets in blood.

These 13 tissues play different roles in virus infection and proliferation. For the viral life cycle, the viruses firstly bind the host receptors on the cell surface. Hence, the distribution of viral receptors in different cell types of diverse tissues can reveal the viral tropism and potential transmission routes. We therefore explored the expression spectrum of host receptors in the following section.

### Expression atlas of ACE2, ssRNA viral receptors and other membrane proteins in 13 human tissues

We firstly analyzed the expression pattern of ACE2 across 13 tissues (**Figure 1**). Our results reveal that ACE2 expresses in lung AT2, liver cholangiocyte, colon colonocytes, esophagus keratinocytes, ileum ECs, rectum ECs, stomach epithelial cells, and kidney proximal tubules, consistent with the recent reports[5–8]. However, ACE2 expression levels are rather low in lung AT2 (4.7-fold lower than the average expression level of all ACE2 expressing cell types). We assume that the presence of co-receptors or other auxiliary membrane proteins in AT2 cells may facilitate the binding and entry of the nCoV.

**Figure 1.**
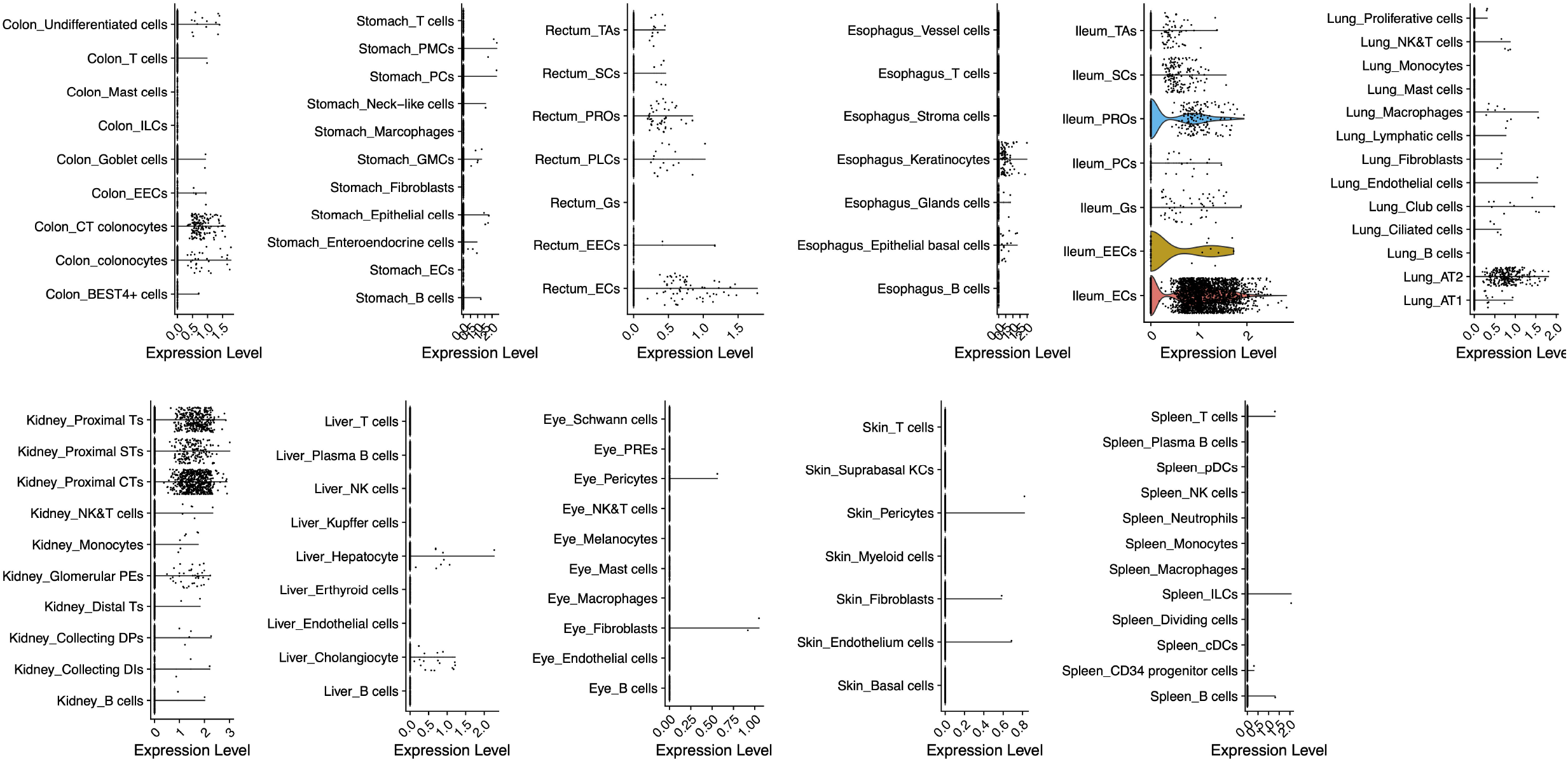
The expression profiles of ACE2 in 13 human tissues. We explored single cell expression maps of ACE2 in lung, liver, stomach, ileum, rectum, colon, blood, bone marrow, spleen, esophagus, kidney, skin and eye, totally comprising 119 cell types (AT1, Alveolar cells Type2; AT2, Alveolar cells Type2; PMCs, pit mucous cells; ECs, endothelial cells; GMCs, antral basal gland mucous cells; PCs, proliferative cells in stomach and Paneth cells in ileum; ECs, enterocytes, EECs, enteroendocrine cells; Gs, goblet cells; PROs, progenitor cells; SCs, stem cells; TAs, transient amplifying; EECs, enteroendocrine cells; RPEs, retinal pigment epithelium). None of the ACE2 transcripts was found in bone marrow and blood.

We then analyzed the co-expression features of the human ssRNA viral receptors and membrane proteins. We collected a total of 152 pairs of virus-host receptor interaction from Viral Receptor database[9] (http://www.computationalbiology.cn:5000/viralReceptor) which curated a number of validated viral receptors. These virus-host receptor interactions contain 51 host receptors in 9 hosts and 96 viruses (**Table S2**). Furthermore, 400 membrane proteins were extracted from Membranome database[25] (https://membranome.org). Totally 451 genes were curated, 95.7% (432/451) of which express in at least one of the 13 tissues. We examined the expression spectrum of the 432 genes across the 119 cell types.

To elaborate the potential relationship between ACE2 and other membrane proteins or viral receptors, we calculated the Pearson Correlation Coefficient between each two genes in the curated reservoir. The findings show that 94 genes are significantly correlated with ACE2 (P<0.01) in a manner of gene expression. Of note, ANPEP, ENPEP and DPP4 are the top three genes correlated with ACE2 (R>0.8) (**Figure 2**). ANPEP, alanyl aminopeptidase, is a host receptor targeted by porcine epidemic diarrhea virus, human coronavirus 229E, feline coronavirus, canine coronavirus, transmissible gastroenteritis virus and infectious bronchitis virus. These viruses all belong to Coronaviridae. Besides skin, ANPEP also mainly expresses in colon, ileum, rectum, kidney and liver (**Figure S3** & **Figure S4**), demonstrating that receptor of coronavirus may have similar expression profiles in human body. ENPEP, Glutamyl Aminopeptidase, belongs to the peptidase M1 family which is the mammalian type II integral membrane zinc-containing endopeptidases. ENPEP regulates blood pressure regulation and blood vessel formation through the catabolic pathway of the renin-angiotensin system[26]. The relationship between ENPEP and viral infection is unknown. DPP4, the receptor of MERS-CoV, shows expression similarity with ACE2, except that DPP4 expresses in some T cells of all the observed tissues (**Figure S3** & **Figure S4**). All of the three genes encode peptidase, which are uniquely adopted by coronavirus as their receptors [27]. This result raised the possibility that ENPEP may be another yet unknown receptor for coronavirus. To further consolidate the findings, we calculated the Euclidean distance between all the curated proteins and constructed their hierarchy relationships across the 119 cell types. DPP4 was the first gene clustered with ACE2.

**Figure 2.**
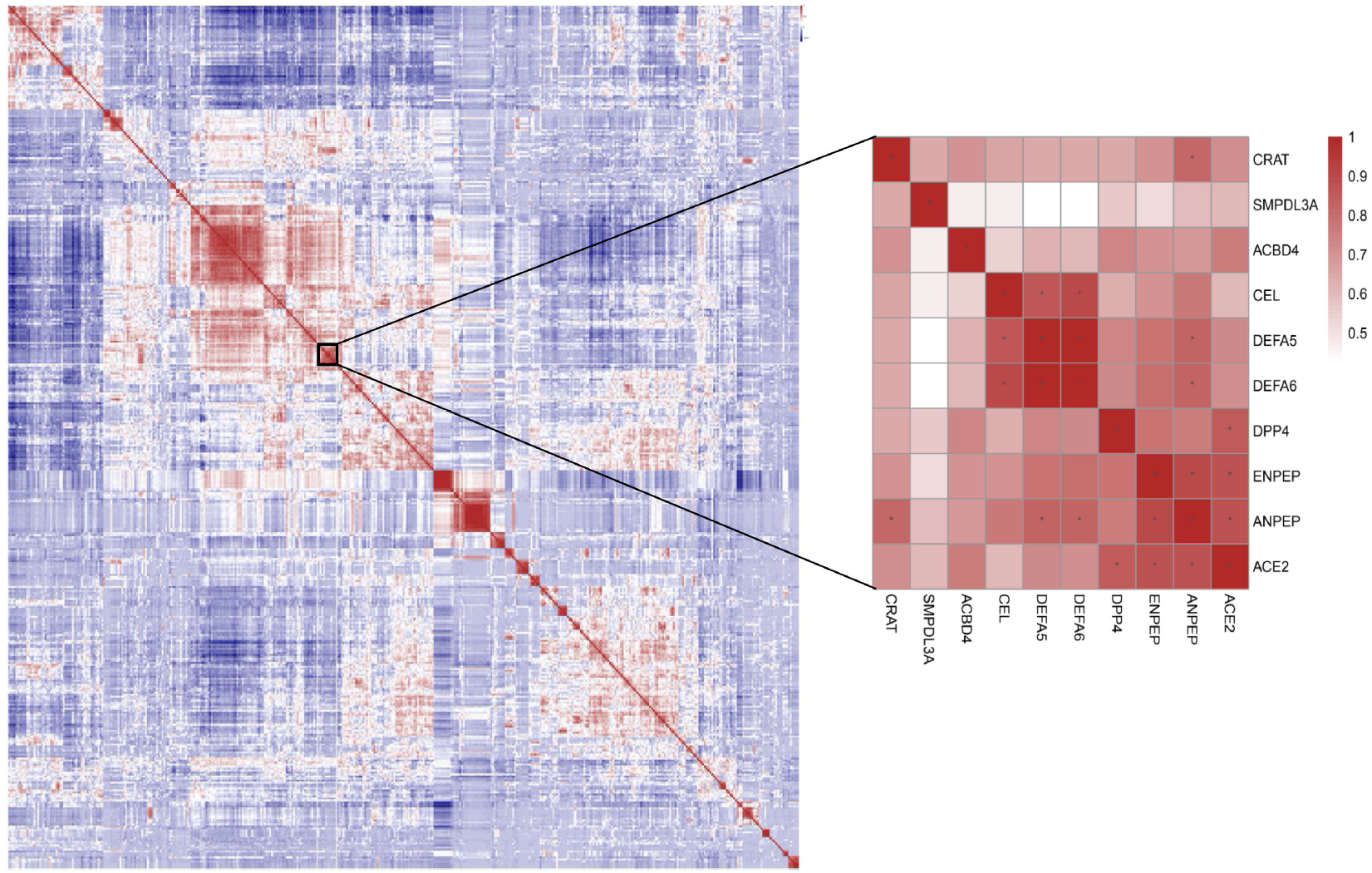
Pearson correlation coefficients between the curated ssRNA viral receptors and membrane proteins. The warm colors mean positive correlation, and the cold colors mean negative correlation. The stars represent the correlation coefficients greater than 0.8. ANPEP, ENPEP and DPP4 show highest correlation with ACE2.

Together, our data demonstrates that the coronavirus receptors tend to share co-expression pattern across different tissues, consistent with the fact that CoVs infect similar types of cells and CoV-infected patients share similar clinical symptoms.

### Macrophages are frequently interacted with the ACE2-expressing cells

Virus-infected cells can recruit and modulate immune cells through secreting chemokines or other cytokines. We sought to identify potential immune cells crosstalking with CoVs-targeted cells. The cell-cell interaction analysis was conducted by CellPhoneDB[24]. The interactions with p-value < 0.01 were adopted to construct the interaction relationship between cell types in each tissue.

Using the cell type expressing ACE2 as ligand-secreting cells, we calculated the total number of interactions with each receptor-secreting cell type. As a result, we found that macrophages showed highest active interaction with ACE2-expressing cells in liver, lung and stomach (**Figure 3A**) sharing a CD74-MIF signaling pairs (**Figure 3B**). CD74 is expressed on the cell surface of antigen-presenting cells and act as a receptor for the cytokine in immune cells. MIF, macrophage migration inhibitory factor, is a pro-inflammatory cytokine participating in inflammatory and immune responses. Besides, PROs, SCs and TAs show high activity responding to the ACE2-expressing cells in ileum and rectum. In colon, ILCs were found frequently interacted with the ACE2-expressing cells. Glomerular parietal epithelial cells and epithelial basal cells in kidney and esophagus also correlated with the cells transcribing ACE2 at very high frequency.

We conclude that the nCoV-targeted cells (ACE2-expressing), can interact with various cell types in different tissues, especially macrophages in lung, liver and stomach. Macrophages may be recruited by nCoV-targeted cells through CD74-MIF interaction and other signaling pathways during infection, play defensive and destructive functions.

**Figure 3.**
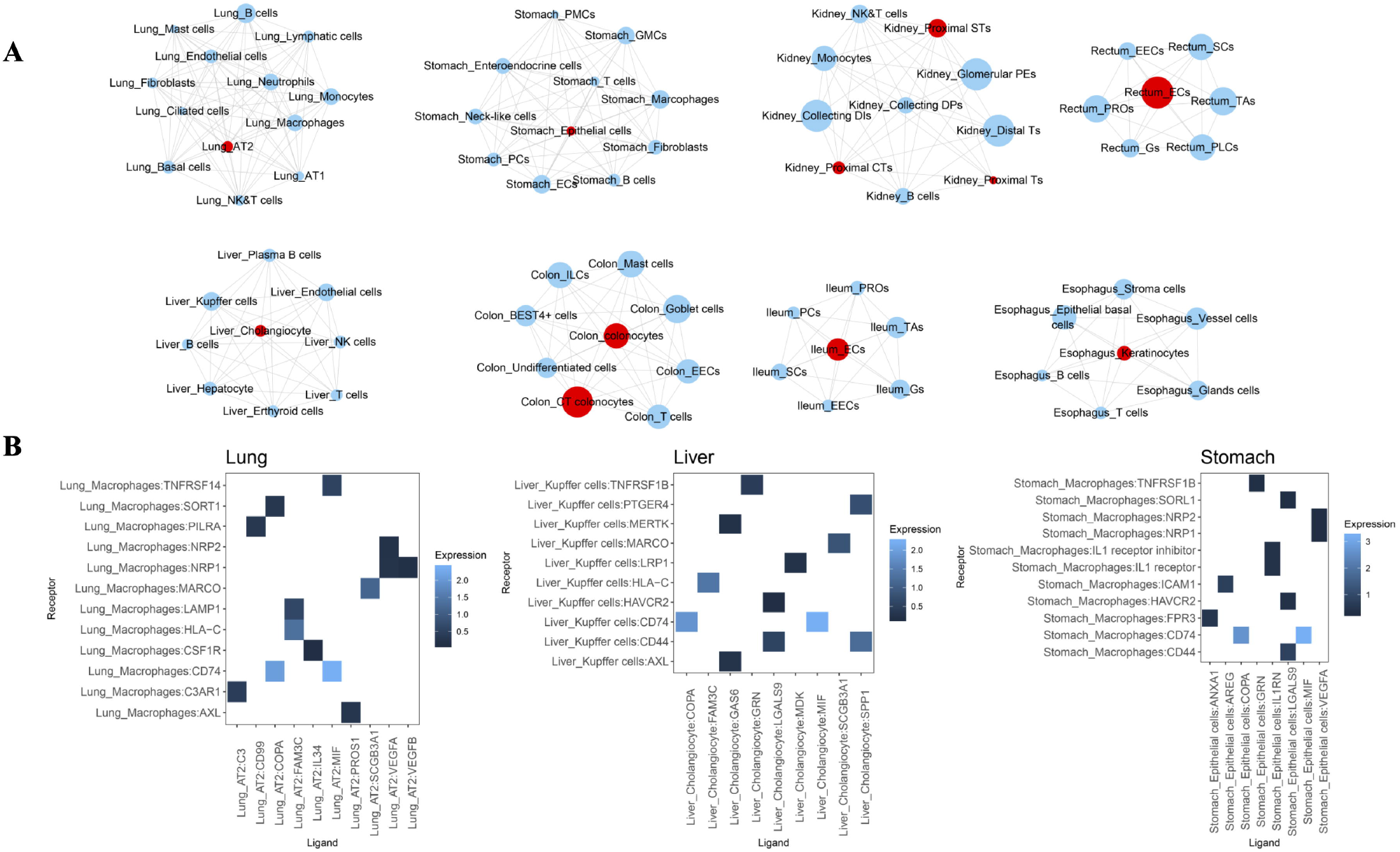
Cell-cell interactions between cell types in 8 tissues expressing ACE2. **(A)** The cell-cell interaction analysis in lung, liver, stomach, ileum, rectum, colon, esophagus and kidney was conducted by CellPhoneDB. Cell types are nodes and interactions are edges. Red nodes indicate major cell types expressing ACE2 in each tissue. Size of cell type is proportional to the total number of interactions with the red nodes. **(B)** The cytokines that connected ACE2-expressing cells and the macrophages. Macrophages expression receptors and ACE2-expressing cells express ligands.

## Discussion

The coronaviruses are a large family of ssRNA viruses causing respiratory diseases in humans. Most of the coronaviruses are associated with mild clinical symptoms, except SARS-CoV and MERS-CoV, showing fatality rate of 9.6% and 34%, respectively[28, 29]. In late December 2019, a novel coronavirus, named 2019-nCoV, emerged in Wuhan, Hubei, China. A total of 37,251 2019-nCoV infected cases involving across different regions of China have been confirmed until February 9, 2020, suggesting that 2019-nCoV is also highly infectious. The phylogenic tree constructed from full-genome sequences indicated that 2019-nCoV is a distinct clade from SARS-CoV and MERS-CoV[4].

The most common symptoms of patients infected with 2019-nCoV are fever and cough[30]. However, a proportion of patients showing multi-organ damage and dysfunction was also observed, including acute respiratory distress syndrome (17%), acute respiratory injury (8%) and acute renal injury (3%). It is also increasingly recognized that 2019-nCoV could be transmitted via multiple routes.

The viruses target host cells via binding host receptors before engaging the infection cycle. ACE2 was proved to be the cell receptor of 2019-nCoV, the same receptor as SARS-CoV[5, 10]. The expression profiles of ACE2 across different cell types of different organs will reveal clues of the virus transmission routes and its potential pathogenesis. In previous studies, ACE2 was found to express in the esophagus upper and stratified epithelial cells, absorptive enterocytes from ileum and colon, alveolar type II cells in lung, liver cholangiocyte and kidney proximal tubules [5–7]. These findings suggested that the clinical symptoms of hepatic failure, respiratory injury, acute kidney injury or diarrhoea may be associated with the pervasive ACE2 expressing cells in these tissues. However, we and others found that ACE2 is lowly expressed, especially in the lung (the main target organ of nCoVs), raising the possible existence of coreceptors facilitating nCoV infection. It is well recognized that ssRNA viruses tend to have multiple receptors[9]. For example, ACE2, CD209 (Dendritic Cell-Specific ICAM-3-Grabbing Non-Integrin 1), CLEC4G (C-type lectin domain family 4 member G) and CLEC4M (C-type lectin domain family 4 member M) are receptors of SARS-CoV[31–34]. In addition, other membrane proteins may also assist virus entry[3]. Since the viral receptor and co-receptors should be co-expressed on the same cell types, we analyzed single cell co-expression patterns covering 400 membrane proteins and 51 known viral receptors in this study. After calculating their gene expression similarity, we found ANPEP, ENPEP and DPP4 are top three genes correlated with ACE4 (R>0.8). Interestingly, both ANPEP and DPP4 are viral receptors of human coronaviruses [35], while ENPEP is also a peptidase, despite that its involvement in virus infection is unclear. For mysterious reasons, human coronaviruses use peptidases as their receptors[27]. Now, we showed co-expression profiles of these molecules, indicating that different human CoVs actually target the similar cell types across different human tissues. It also explains why, patients infected with different human CoVs manifest similar clinical symptoms. We propose that further experimental validations should be performed to explore the role of these peptidase in 2019-nCoV and other CoVs infection.

Host immune response plays crucial roles in the fight against viruses. Generally, virally infected cells release interferons to suppress viral activities [36–38]. The interferons also act on warning the neighboring cells of virus attack. It can signify the the nearby cells to upregulate MHC class I molecules to notify the CD8^+^ T cells to identify and eliminate the viral infection[39, 40]. Understand the potential cell-cell communication mechanisms across different tissues is important for understanding immune reactions. In this study, we investigated the cells communicated with CoVs-targets (ACE2-expressing cells) in each tissue. Our results illustrate that macrophages frequently crosstalk with the ACE2-expressing cells, in lung, liver and stomach etc. This suggests that macrophages play the sentinel role during human CoVs infection. Future studies should investigate these signaling pairs in the setting of CoVs infection in patients and animal models.

---

**Figure S1 Cell type identification in 13 human tissues.** The cell types were annotated by curated known cell markers searched from their corresponding publications (see **Table S1** and materials and methods).

**Figure S2 Cell type proportion for each human tissue.** Proportion of each cell type was calculated for each tissue.

**Figure S3 Heatmap of ANPEP, ENPEP and DPP4 expression in 13 human tissues.** We depicted single cell expression maps of ANPEP, ENPEP and DPP4 in lung, liver, stomach, ileum, rectum, colon, blood, bone marrow, spleen, esophagus, kidney, skin and eye, totally including 119 cell types. The expression was were log scaled.

**Figure S4 Violin plot of ANPEP, ENPEP and DPP4 expression in 13 human tissues**

