## Supplementary figures and images for "Single cell RNA sequencing of 13 human tissues identify cell types and receptors of human coronaviruses"

### Figure S1

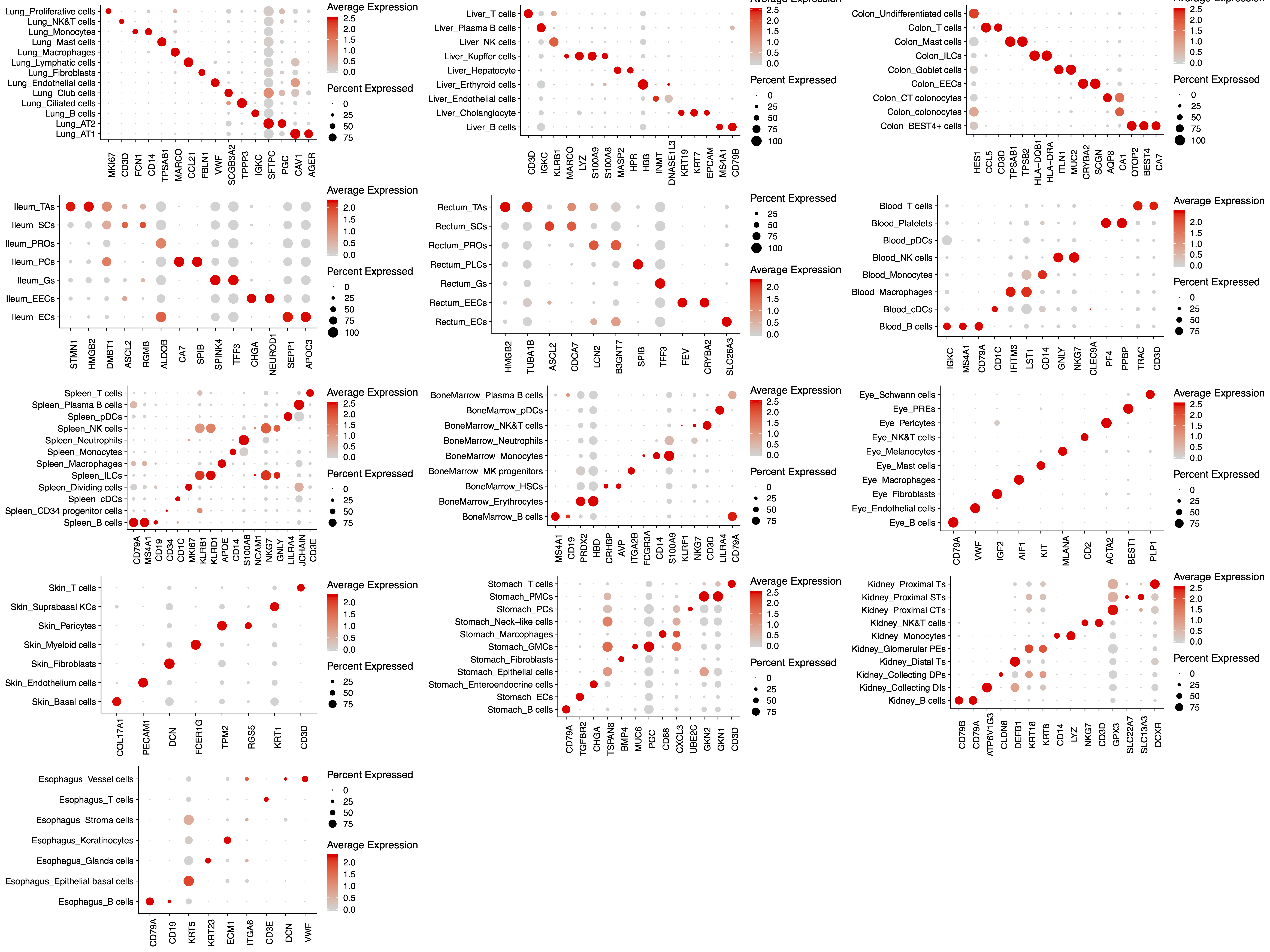

### Figure S2

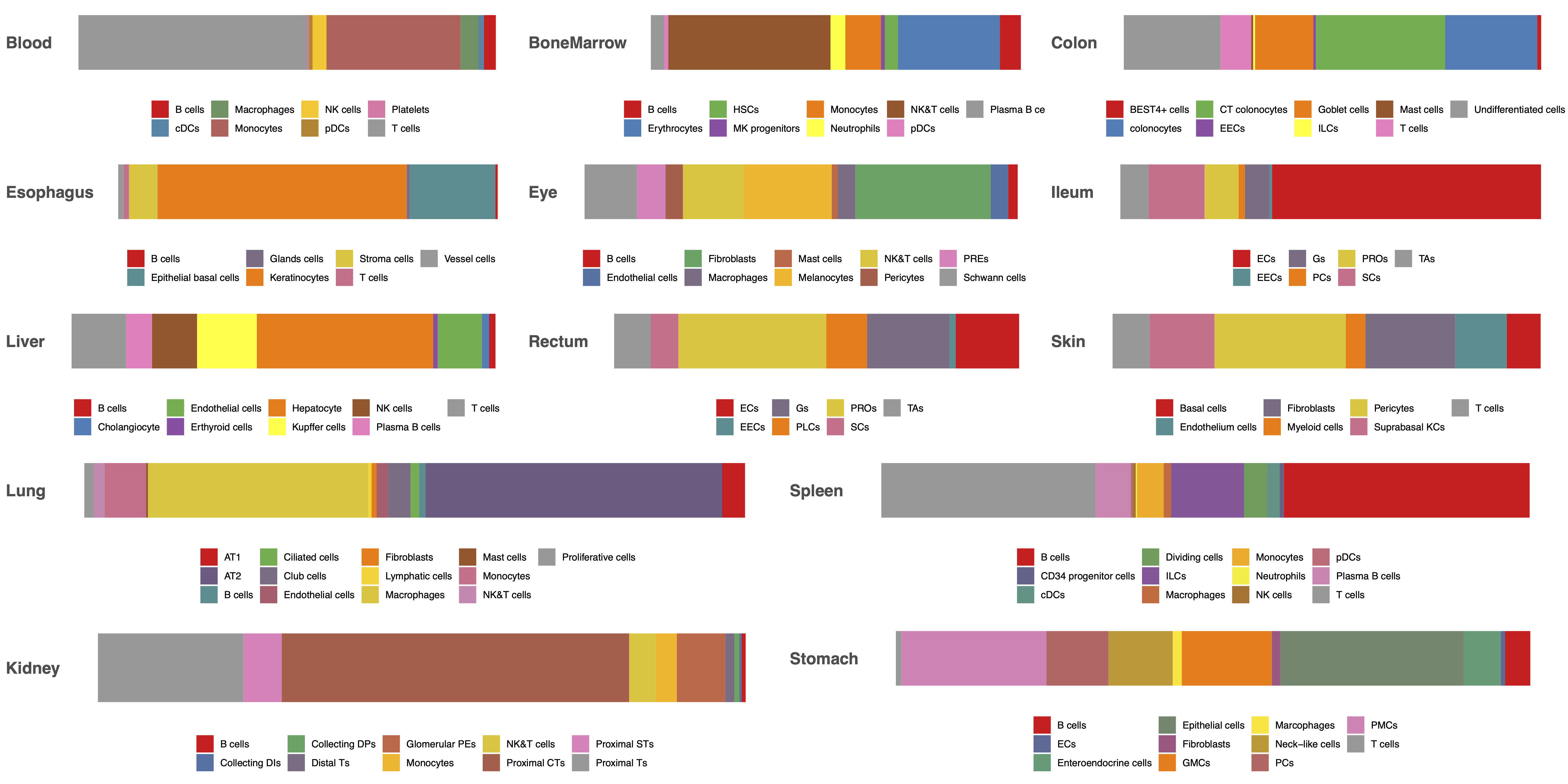

### Figure S3

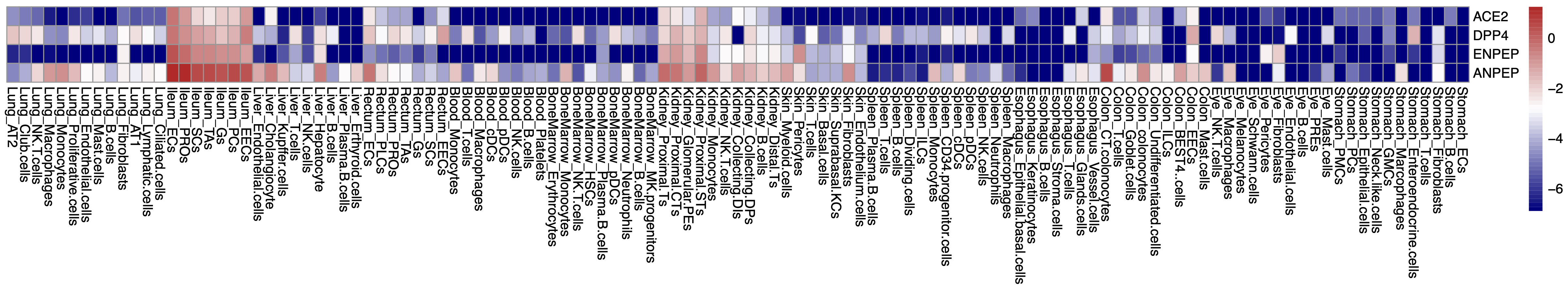

### Figure S4

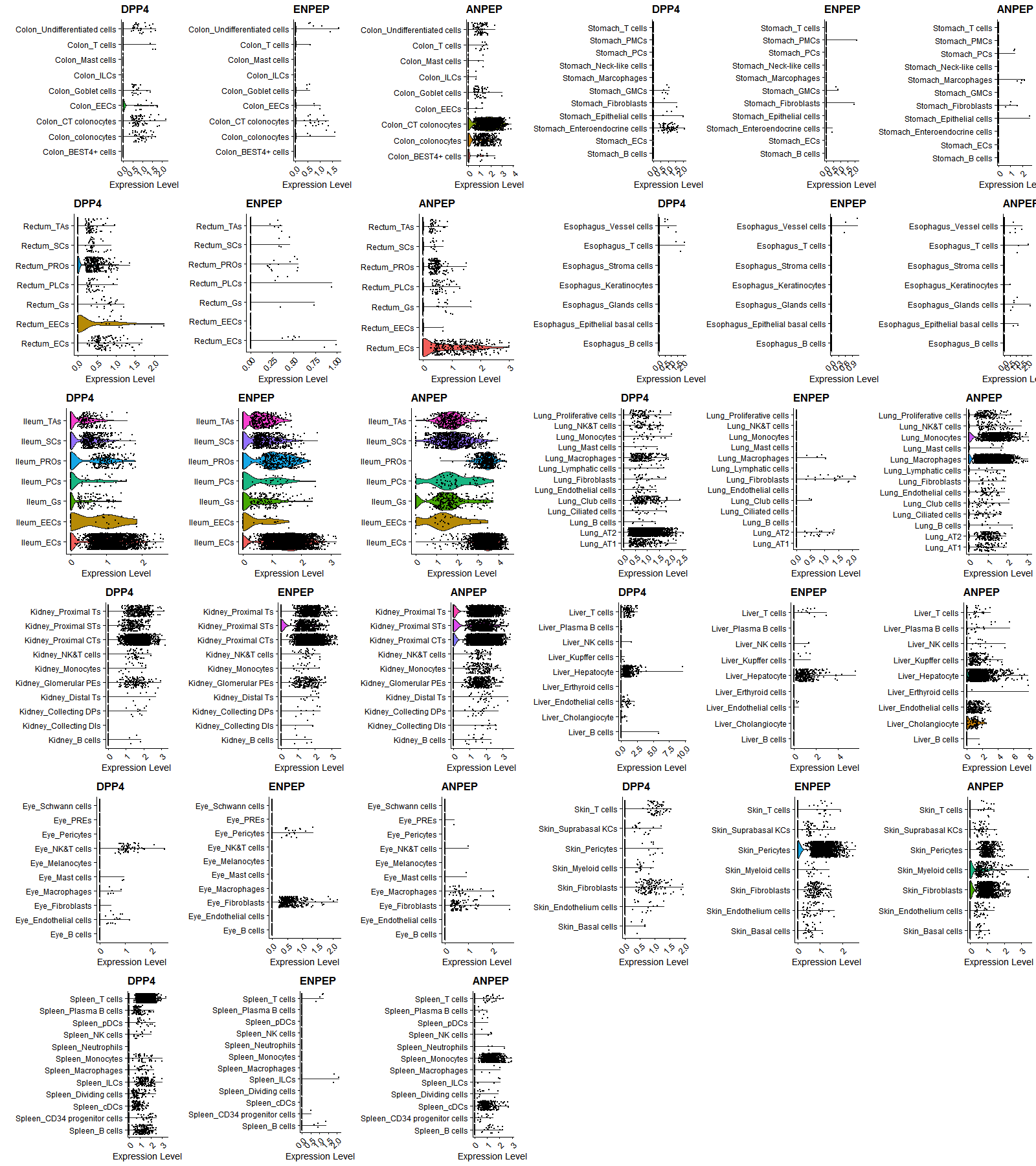
